# CIT tumor lines: A novel series of immunogenic cutaneous squamous cell carcinoma cell lines derived from chemical carcinogenesis

**DOI:** 10.1101/2025.04.03.647071

**Authors:** Robert Letchworth, Miho Tanaka, Alina A Barnes, Lotus Lum, Savannah Hughes, Grant Schlauderaff, Piyush Chaudhary, Kenneth M Ng, Daphne Superville, Corinna Martinez Luna, Maria Gonzalez, Dekker C Deacon, Allie Grossmann, Melissa Q Reeves

**Affiliations:** Huntsman Cancer Institute, University of Utah; Department of Pathology, University of Utah; Department of Microbiology and Immunology, University of California, San Francisco; Department of Dermatology, University of Utah; Earle A. Chiles Research Institute, Providence Cancer Institute of Oregon, Portland, OR

**Keywords:** Cutaneous squamous cell carcinoma, tumor immunology, tumor neoantigen, immune checkpoint inhibitor, mouse models of cancer

## Abstract

Immunotherapy is now widely used to treat advanced-stage skin cancer, but it is effective for only approximately half of skin cancer patients, including both patients with melanoma and cutaneous squamous cell carcinoma (cSCC). To overcome current barriers, preclinical mouse models that faithfully recapitulate the tumor genetics, mutation burdens, and neoantigen patterns of specific human tumor types are essential. However, while many models exist for melanoma, there are currently relatively few transplantable murine models of cSCC, which is responsible for nearly as many deaths as melanoma each year. Here we describe a novel series of 11 cSCC tumor lines, the Carcinogen-Induced Tumor (CIT) lines, syngeneic to the FVB strain, that address this need. The CIT lines were established from skin carcinomas induced by DMBA and TPA treatment and harbor genetic drivers and overall tumor mutational burdens that recapitulate those found in cSCC. Each CIT line gives rise to tumors with a consistent immune infiltration pattern, ranging from T cell-rich “hot” tumors to T cell-poor “cold” tumors. Hot CIT lines exhibit partial responses to treatment with immune checkpoint inhibitors, and we have identified two neoantigens present in an immunotherapy-responsive CIT line. The CIT lines thus provide a valuable new series of preclinical models for studying anti-tumor immune responses and developing strategies to improve immunotherapy efficacy in cSCC.

## Background

Cutaneous squamous cell carcinoma (cSCC) is the second most common cancer type in the United States, affecting approximately 700,000 people annually^1–3^ and responsible for between 8,000 and 15,000 deaths per year^1,4^. Although cSCC carries a relatively low risk of local advancement or distant metastasis, its high incidence rate means that as many or more Americans are estimated to die every year from cSCC as from melanoma^4^. Among high-risk cSCC patients, up to 5% will progress to locally advanced or metastatic tumors that cannot be treated with surgery or radiation alone^1,5,6^. Immune checkpoint inhibitors (ICIs), which have revolutionized cancer treatment over the past decade^7,8^, are approved for advanced cSCC, and hold immense potential to be curative for these patients^9,10^. However, overall response rates of cSCC to ICI therapy are only 50-60% in locally advanced disease^10–12^ and drop as low as 35% in metastatic disease^11^. These statistics highlight an unmet clinical need to understand why ICI fails for many cSCC patients and how immunotherapy strategies can be improved for these patients^10–13^.

Murine models play a vital role in basic and translational cancer research, providing a means of conducting mechanistic experiments to dissect cancer and immune cell biology and their interactions with therapy. Transplantable tumor models, in which cancer cell lines are implanted into immune-competent syngeneic hosts, play a particularly important role in preclinical research, as the reliability of their growth kinetics makes them a workhorse for testing the efficacy of new therapies or combinations. Additionally, transplantable murine models are essential for studying immunotherapy. Because ICIs primarily act directly on T cells, they are effective only in immune-competent hosts, making patient-derived xenografts (PDXs)—which grow only in mice with compromised immune systems—unsuitable models to study immunotherapy. However, while transplantable murine models of melanoma are widely available and have contributed to substantial advancements in immunotherapy, relatively few transplantable tumor models are available for cSCC. Many barriers to the efficacy of immunotherapy are cancer type-specific, and there is a need for mouse models that accurately recapitulate the unique features of each disease. As such, the lack of availability of transplantable cSCC models represents a significant gap in the field.

Here we describe the Carcinogen-Induced Tumor (CIT) cell line panel, a novel series of murine cSCC cell lines, to address this need. The CIT lines are derived from DMBA/TPA-induced cutaneous SCCs, and they are syngeneic to the FVB strain. Notably, the CIT cell lines recapitulate the mutational profiles of human cSCC, including harboring cSCC-relevant driver mutations in *Hras* and *Trp53* and deletions of *Cdkn2a*. As a result of being chemically induced in immune-competent hosts, the CIT lines carry a moderate to high mutation burden, like human cSCC^14,15^. The tumors that arise from the CIT lines exhibit a range of reproducible patterns of immune infiltration—from high to low T cell infiltrate—along with a corresponding range of responses to immunotherapy. Additionally, as a consequence of their high mutation burden, the CIT lines harbor endogeneous neoantigens, which are mutant proteins—resulting from genetic mutations—that can be recognized by T cells^8^. Neoantigens enable T cells to identify tumor cells as “foreign,” and this recognition is a critical central mechanism of the anti-tumor immune response that is enhanced by ICI therapy^8,16^. The presence of these neoantigens—two of which we describe in this manuscript—are an additional valuable tool to facilitate the study of tumor-specific T cells during immunotherapy responses in cSCC.

The CIT cell line series thus offers a panel of cSCC tumor models that exhibit a range of mutational burdens, immune infiltrates, and neoantigens, while being driven by cSCC-relevant genomic drivers and exhibiting histopathologic features characteristic of cSCC. These properties make the CIT lines a valuable set of preclinical models for investigating tumor biology and responses to therapy, particularly immunotherapy, in cSCC.

## Results

### CIT lines display a range of reproducible immune infiltration phenotypes

The CIT cell lines were derived from skin carcinomas that arose by topically treating *K5-CreER^T2^*-*Confetti* FVB/N mice with a single dose of 7,12-dimethylbenzanthracene (DMBA) followed by 12-O-tetradecanoylphorbol-13-acetate (TPA) twice weekly for 20 weeks, using a well-established skin carcinogenesis protocol^17,18^ (**Fig. 1A**). In the DMBA/TPA model, benign papillomas arise after about 6 weeks of TPA treatment, and approximately 10% of these progress to malignant carcinomas. Primary carcinomas were resected when they were approximately 1cm in longest diameter, digested into a single cell suspension, and grown under standard tissue culture conditions. Each cell line was established in tissue culture for >10 passages. Implantation of 1.25×10^5^ tumor cells into the subcutaneous flank of FVB mice gave rise to tumors with 100% penetrance, and we subsequently profiled the histopathologic and immune characteristics of these tumors.

**Figure 1.**
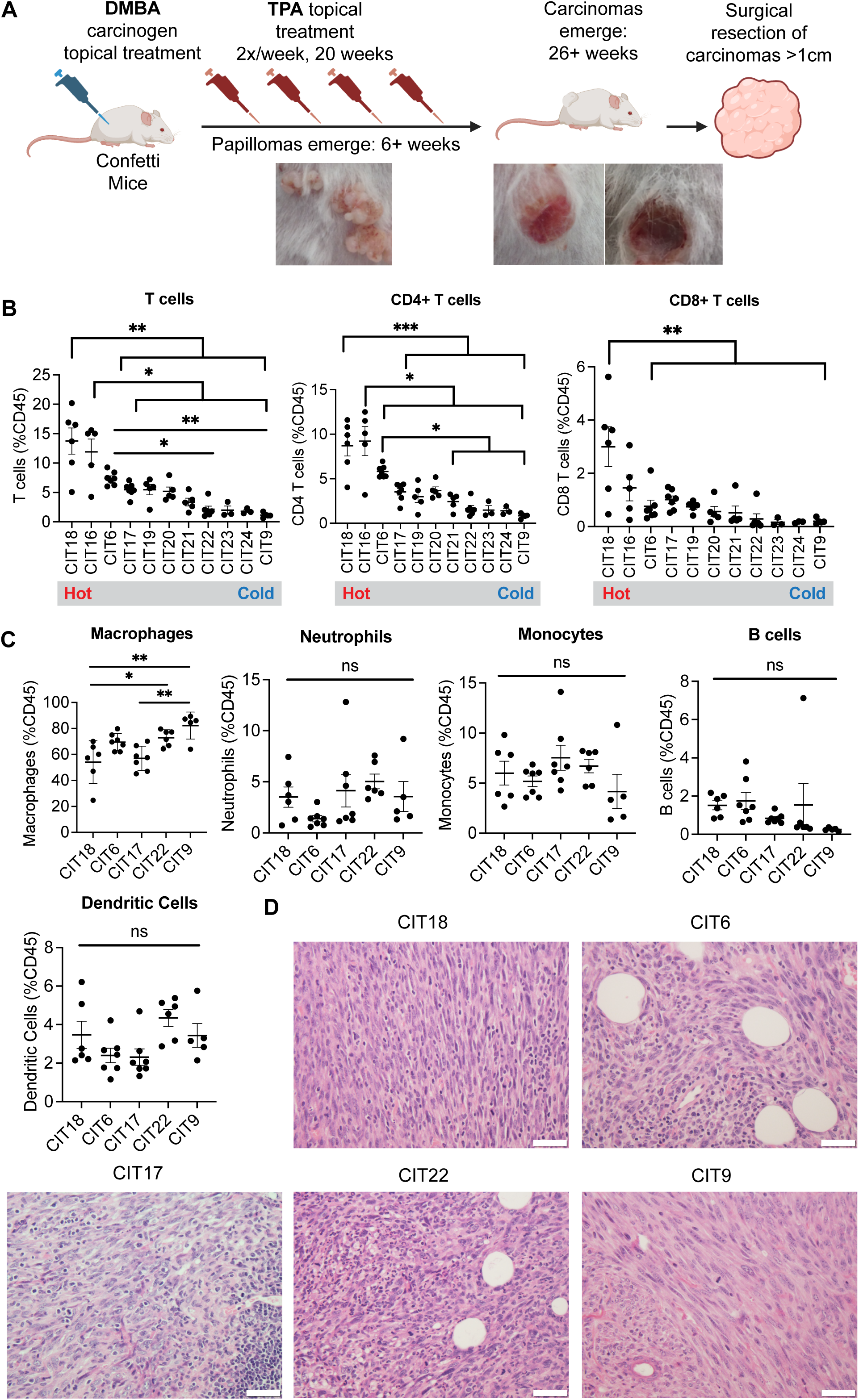
Immune infiltration and histology profiles of CIT lines. **(A)** Schematic of CIT cell line generation. Tumors were induced with the DMBA/TPA carcinogenesis model, and cell lines were established from malignant carcinomas. **(B)** Quantification of total T cells, CD4+ T cells and CD8+ T cells by flow cytometry in each CIT line. (n=5-7 mice per group; * *p* < 0.05, ** *p* < 0.01, *** *p* < 0.001) **(C)** Quantification of macrophages, neutrophils, monocytes, B cells and dendritic cells by flow cytometry in 5 selected CIT lines. (n=5-7 mice per group; * *p* < 0.05, ** *p* < 0.01, ns = not significant) **(D)** Representative images of tumors from selected CIT lines stained with hematoxylin and eosin (H&E) for histological analysis. Scale bar = 20 μm. Statistical analysis in (B-C) by one-way ANOVA with Tukey post-hoc test.

We carried out immune profiling of 11 CIT cell lines by flow cytometry, quantifying the number of total T cells, CD4+ and CD8+ T cells in tumors from each CIT model (**Supp. Fig. 1A**). We found each CIT line gave rise to tumors with a consistent pattern of immune infiltration, spanning the range of immune “hot” (highly infiltrated by T cells, e.g., CIT18) to immune “cold” (poorly infiltrated by T cells, e.g., CIT9) (**Fig. 1B**). Interestingly, we observed a consistent bias in the T cell compartment toward CD4+ T cells, about half of which were Tregs (**Supp. Fig. 1B**). We selected 5 CIT lines, including representation of both hot and cold models, for more extensive immune profiling. On the whole, we found the myeloid compartment of CIT tumors was strongly biased toward macrophages (CD11b+MHCII+F4/80+), with relatively fewer Ly6C+ and Ly6G+ myeloid cells (monocytes and neutrophils, respectively) and DCs (CD11c+MHCII+) (**Fig. 1C, Supp. Fig. 1A**). Macrophage infiltration was highest in the two coldest (lowest T cell infiltrate) CIT models, CIT22 and CIT9. Conversely, hot models CIT18 and CIT6 exhibited the highest B cell infiltration (**Fig. 1C**). There was no clear trend between hot and cold tumors for Ly6G^hi^Ly6C^mid^ neutrophils, Ly6C^hi^CD11b+ monocytes, or CD11c+MHCII+ dendritic cells (**Fig. 1C**).

Tumor histology for the same five selected CIT tumor models was reviewed by a board-certified pathologist. Tumors from all five CIT models were poorly differentiated, spindled, arranged in fascicles, and highly mitotic (**Fig. 1D**). Some CIT18 tumors contained pleomorphic, rhabdoid, or epithelioid malignant cell morphologies. Among the hot models, CIT18 and CIT17 tumors contained brisk tumor-infiltrating lymphocyte (TIL) infiltration, while CIT6 tumors contained focal clusters of TILs or TILs localized to the edge of the tumor. In comparison, tumors arising from the cold lines CIT9 and CIT22 displayed very little TIL infiltration. CIT6, CIT17, CIT18 and CIT22 tumors contained diffuse infiltration of polymorphonuclear (PMN) cells, while PMNs in cold CIT9 tumors were largely restricted to the edge of the tumors. Altogether, we find that the CIT cell lines give rise to tumors that recapitulate the histologic features of high grade, sarcomatoid SCC, and each exhibits characteristic patterns of immune infiltration congruent with flow cytometry-based immune profiling.

### Genomic profiles of CIT lines recapitulate human SCC

All 11 CIT cell lines were subjected to exome sequencing, to a median depth of 153X. Consistent with previous sequencing of DMBA/TPA tumors by ourselves and others^17,19,20^, CIT cell lines were genomically characterized by *Hras* driver mutations, *Cdkn2a* losses, a substantial mutation burden, and whole-chromosome copy number alterations. The mutation burden among the CIT lines ranged from 174 to 703 mutations (5.2-21.0 mutations/Mb; average of 10 mutations/Mb), including 46 to 256 non-synonymous mutations (**Fig. 2A-B, Supp. Table 1**). Ten of our CIT lines exhibited an *Hras* Q61L mutation, while one, CIT16, exhibited an *Hras* G12V as well as a *Kras* G13R mutation (**Fig. 2C**). Seven CIT lines displayed additional non-synonymous mutations in driver genes known to be mutated in human cSCC, including *Trp53* and *Fgfr3*, while nine CIT lines harbored a deletion at the *Cdkn2a* locus (**Fig. 2C**). The majority of total mutations were T>A transversions, with a particular prevalence of CTN>A mutations (with N representing any nucleotide) consistent with the known mutagenic signature of DMBA^17,19,20^ **(Fig. 2D)**. While this signature is not identical to the C>T mutations induced by UV and typically found in cSCC^14,15^, we note that the genes mutated in the CIT lines were nonetheless cSCC-relevant driver genes, including mutations at classical *Hras* hotspots in codons 12 and 61. The CIT lines also harbored whole-chromosome alterations, primarily amplifications, especially on chromosomes 6 and 15 **(Fig. 2E)**. Chromosome 6 amplifications are common in DMBA/TPA-induced skin carcinomas^17^. Chromosome 15 amplifications were also previously observed, but they were more heavily represented in the CIT lines than in a previously sequenced cohort of primary tumors^17^, suggesting perhaps a selection in cell culture for lines with this amplification. Interestingly, we saw only one whole-chromosome amplification each of chromosome 1 and chromosome 7. We previously reported that chromosome 1 and 7 amplifications are common in DMBA/TPA carcinomas with an epithelial-like “squamous” morphology, but largely absent in DMBA/TPA carcinomas with a mesenchymal-like “spindle” morphology^17^. The underrepresentation of these alterations is perhaps not surprising given that the CIT lines primarily give rise to tumors with spindle cell morphology, but may again suggest a selection bias of the cell lines we were able to establish in culture.

**Figure 2.**
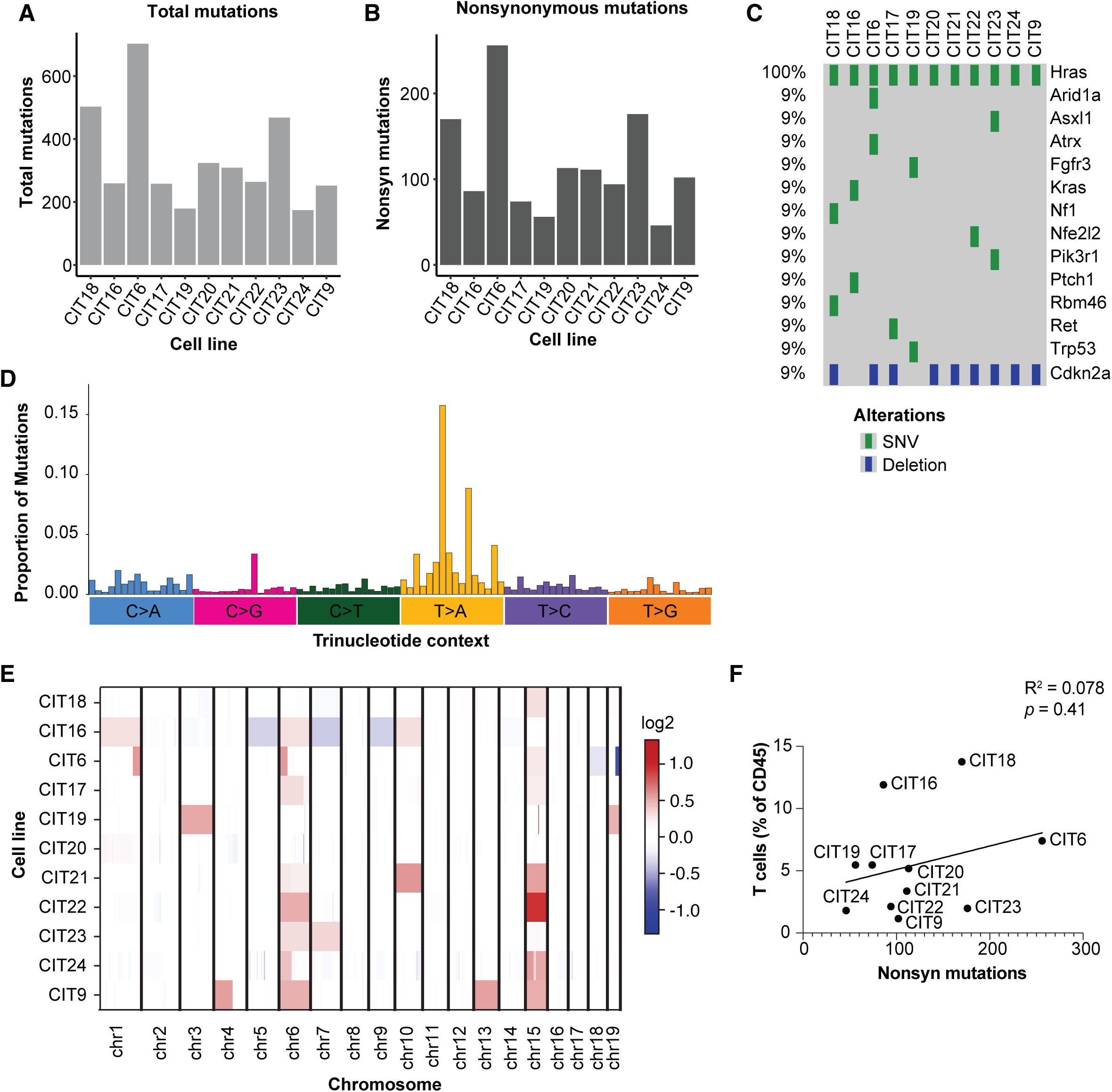
Genetic profiling of CIT lines. **(A-B)** Number of total (A) and non-synonymous (B) mutations per cell line, based on exome sequencing. **(C)** Genes mutated in CIT lines which are known to be commonly mutated in human SCC, and CIT lines with *Cdkn2a* deletions. **(D)** Frequencies of mutations observed in each of 96 possible trinucleotide contexts for all mutations in CIT cell lines. Trinucleotide contexts, arranged on the x axis, are grouped by the base pair change of the mutation. Highest peaks are observed at CTG>A and GTG>A. **(E)** CNV alterations detected in each chromosome in CIT lines based on exome sequencing. Red denotes copy number gains, blue denotes copy number losses. **(F)** Scatter plot showing relationship between non-synonymous mutations (based on exome sequencing) and T cell infiltrate (based on flow cytometry) for CIT lines. There is no significant correlation between these metrics.

Given that mutation burden information and immune phenotypes were available for all 11 cell lines, we examined whether a relationship existed between these metrics. While two of the tumors with the highest T cell infiltration, CIT18 and CIT6, exhibited two of the three highest mutation burdens, we observed that tumor model with the second-highest T cell infiltration, CIT16, had a relatively low mutation burden. CIT23 tumors, which had the second-highest mutation burden, were by contrast very poorly infiltrated by T cells. Thus, aligning with observations from patients, mutation burden alone was insufficient to predict the immune response to any given tumor, and there was not a significant correlation between mutation burden and T cell infiltrate across CIT models (**Fig. 2F**).

### Immunologically hot CIT lines respond to immunotherapy

Because our CIT lines exhibited moderate to high tumor mutation burdens, we hypothesized they would carry neoantigens and would be sensitive to immune checkpoint inhibitors (ICIs). We tested the efficacy of a combination ICI regimen—anti-PD-1 plus anti-CTLA-4, administered as two doses 3 days apart—on CIT18, CIT6, and CIT9 tumors (**Fig. 3A**).

**Figure 3.**
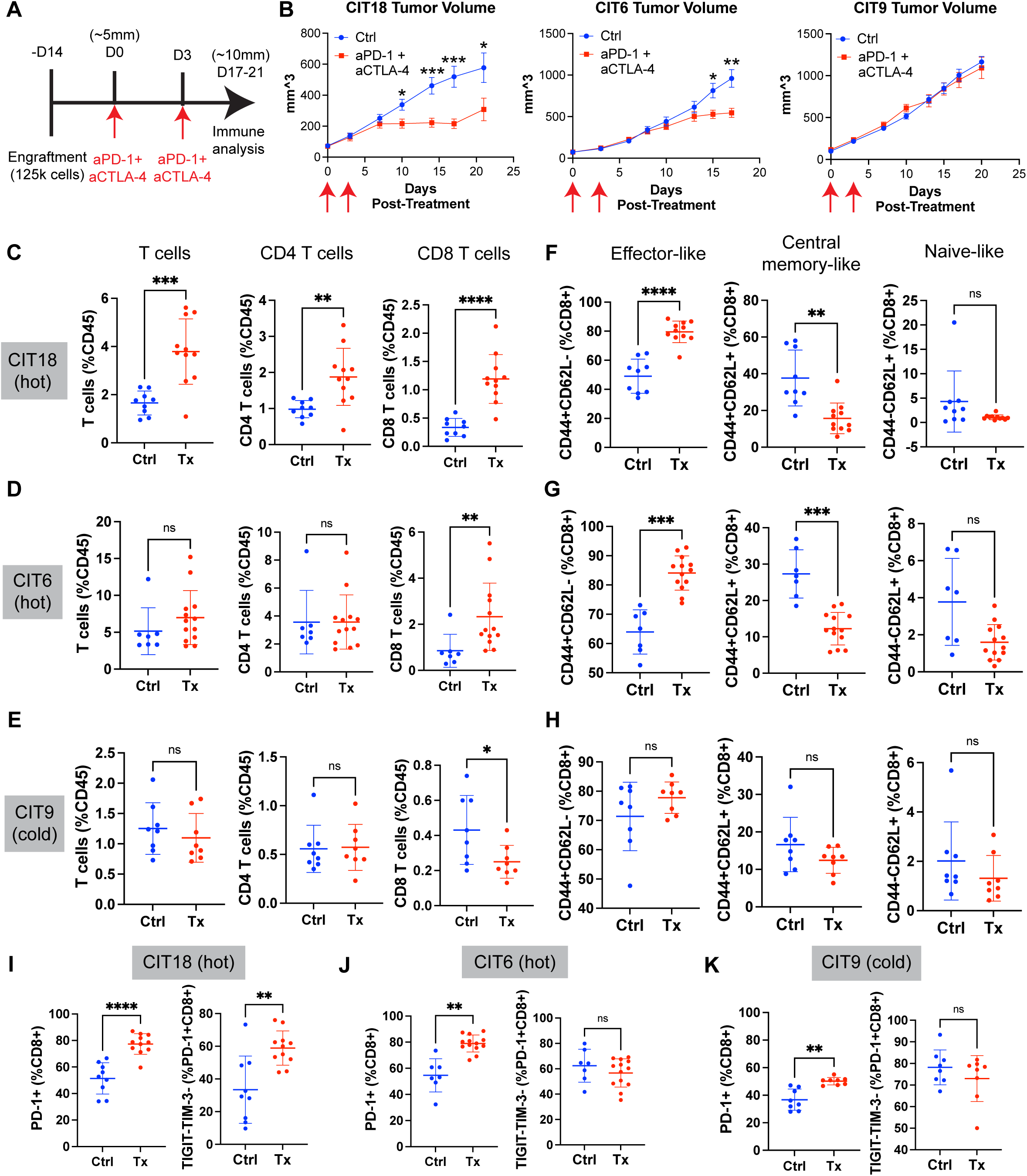
Immunologically hot CIT lines are responsive to combined anti-PD-1 and anti-CTLA-4 immune checkpoint inhibitor (ICI) therapy. **(A)** Schematic of ICI treatment regimen on CIT tumor-bearing mice. **(B)** Average tumor volumes of immunologically hot CIT18 and CIT6 tumors and immunologically cold CIT9 tumors after combined ICI therapy, compared to tumors treated with isotype control antibodies. Error bars represent mean and standard error. **(C-E)** Total T cell, CD4+ T cell, and CD8+ T cell infiltration of CIT18 (C), CIT6 (D) and CIT9 (E) tumors as a percentage of all CD45+ immune cells for control-treated (“Ctrl”) or ICI-treated (“Tx”) tumors. **(F-H)** Abundance of effector subsets of CD8+ T cells defined by the combinatorial expressions of CD44 and CD62L in CIT18 (F), CIT6 (G) and CIT9 (H) tumors. **(I-K)** Abundance of CD8+ T cells expressing PD-1, and frequency of “activated” PD-1+Tim3-TIGIT-CD8+ T cells in CIT18 (I), CIT6 (J) and CIT9 (K) tumors. All statistical analyses performed with Welch’s t-test, n=7-13 mice per group. * *p* < 0.05, ** *p* < 0.01, *** *p* < 0.001, **** *p* < 0.0001.

CIT18 and CIT6 tumors—both of which we had observed to be immunologically hot— grew significantly slower than controls when treated with anti-PD-1 plus anti-CTLA-4 (hereafter, “ICI”) **(Fig. 3B)**. Conversely, immunologically cold CIT9 tumors displayed no difference in growth between ICI-treated and control tumors **(Fig. 3B)**. We carried out immune profiling of ICI- and control-treated tumors to interrogate how ICI remodeled the immune response in these models. CIT18 tumors had the strongest response to ICIs, both in terms of the reduction in growth and in the increase in tumor infiltrating T cells. Total T cells, CD8+ T cells and CD4+ T cells were all more abundant in ICI-treated CIT18 tumors (**Fig. 3C**, **Supp. Fig. 2A**). CIT6 tumors, which showed a more modest response to ICI than CIT18, only exhibited an influx of CD8+ T cells following ICI, while CIT9 tumors showed no increase in overall T cell infiltration. **(Fig. 3D-E**, **Supp. Fig. 2A)**.

ICI treatment also altered the activation profile of tumor-infiltrating T cells. Effective ICI treatment in CIT18 and CIT6 tumors was accompanied by an increase in the percentage of CD8+ and CD4+ T cells which exhibited a CD44+CD62L- “effector-like” phenotype **(Fig. 3F-G**, **Supp. Fig. 2B).** Notably, in hot tumors this increase in “effector-like” CD8+ T cells resulted in a skewing of the CD8+ compartment away from a CD44+CD62L+ “central memory-like” phenotype, even though the abundance of central memory-like CD8+ T cells as a percent of all immune cells was unchanged **(Supp. Fig. 2C)**. In contrast, ICI treatment in cold CIT9 tumors was unsuccessful at shifting the phenotype of either CD8+ or CD4+ T cells **(Fig. 3H**, **Supp. Fig. 2B-C)**. We further examined the expression of PD-1, which is present on both activated and exhausted T cells, and Tim3 and TIGIT, both of which mark exhaustion, on CD8+ T cells. The proportion of tumor-infiltrating CD8+ T cells expressing PD-1 increased after ICI treatment in all three tumor models **(Fig. 3I-K)**. Interestingly, the “activated” PD-1+Tim3-TIGIT- T cell population was the primary CD8+ population to increase in CIT18 tumors following ICI **(Fig. 3I, Supp. Fig. 2D)**, and this corresponded to the best tumor growth control. By contrast in CIT6 tumors, which showed more modest tumor growth control, the proportion of “activated” PD-1+Tim3-TIGIT- CD8+ T cells did not increase as a fraction of PD-1+ CD8+ T cells **(Fig. 3J)**. Instead, there was an increase in the proportion of PD-1+ CD8+ T cells with an exhausted (PD-1+Tim3+ or PD-1+Tim3+TIGIT+) phenotype after ICI treatment **(Supp. Fig. 2D)**. Together, this data shows that CIT tumor models exhibit a range of responses to immunotherapy, and that the degree of sensitivity appears to be approximately correlated with the level of T cell infiltration in untreated tumors.

### CIT tumors display identifiable and immunotherapy-sensitive neoantigens

Cytotoxic CD8+ T cells recognize tumor cells by using their T cell receptor (TCR) to detect neoantigens, which are fragments of mutant proteins that arise from genetic mutations in the tumor^8,16^, and which allow T cells to see tumor cells as “foreign.” Given that improvement in the CD8+ T cell response corresponded to effective tumor control following ICI, we next sought to identify the neoantigens recognized by CD8+ T cells during this response. We focused here on the CIT6 tumor model, which exhibited the intermediate response to ICI among the three models tested and thus may be most representative of patient tumors which are recognized by the host immune response but benefit only marginally from current ICI strategies. We employed the NetH2pan algorithm^21^ to predict the binding strength between the H2-D^q^ and H2- K^q^ MHCI alleles carried by FVB/N mice and the neoepitopes predicted to be generated by all non-synonymous mutations identified in our exome sequencing analysis. We identified 27 top binders, and these were further filtered to prioritize those for which the mutant sequence was predicted to exhibit better binding than the wild-type sequence, and for which we could detect expression of the gene by RNAseq (**Fig. 4A-B**, **Supp. Table 2**).

**Figure 4.**
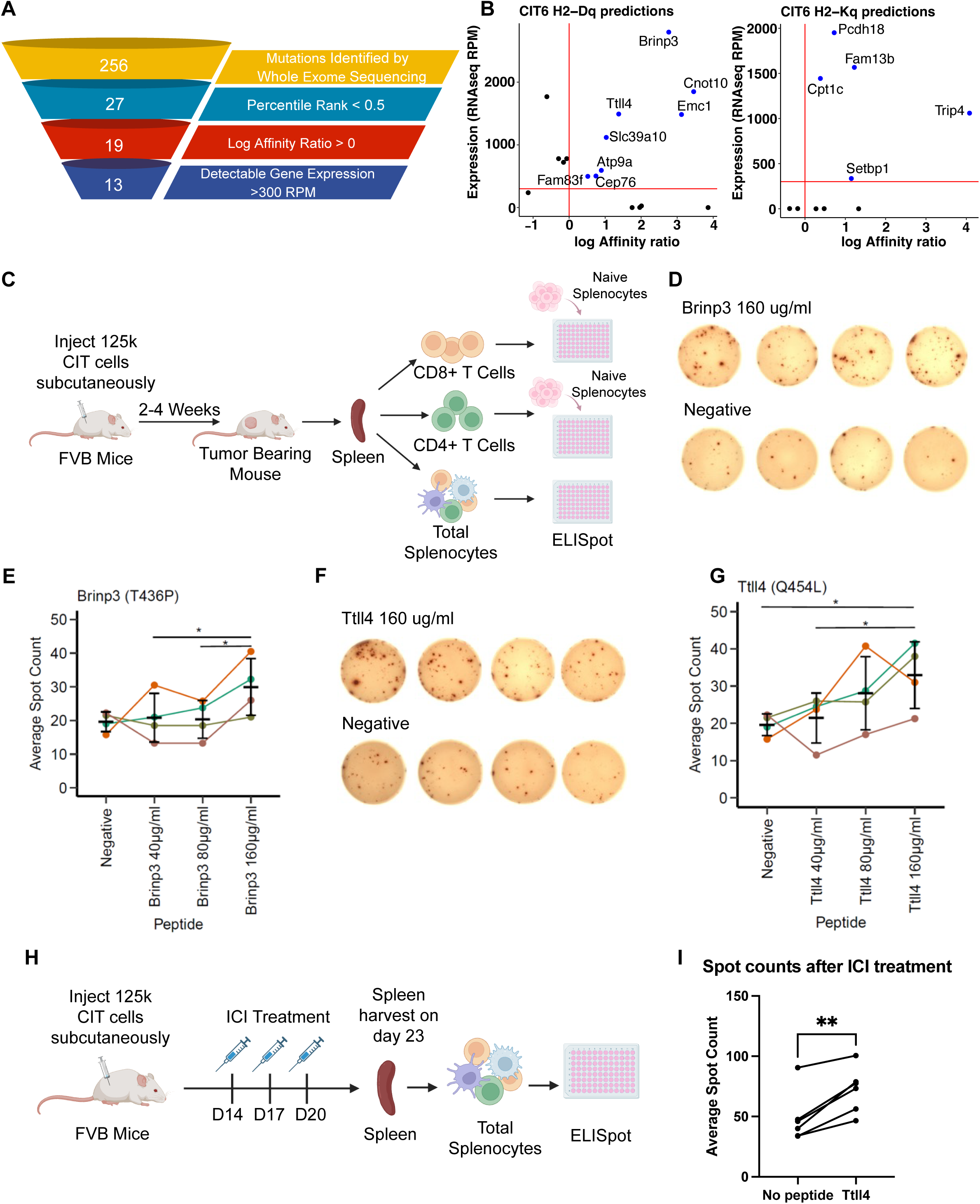
CIT lines express identifiable neoantigens whose cognate CD8+ T cells are responsive to immunotherapy. **(A)** Selection criteria for CIT6 mutations predicted to generate neoantigens. **(B)** Predicted peptide log affinity ratio (affinity ratio is defined as ratio between predicted binding affinity of mutant versus wild-type sequence) and gene expression level of mutations predicted to be top CIT6 candidate neoantigens, shown for H2-Dq and H2-Kq alleles. Affinity predictions were generated using NetH2Pan algorithm. **(C)** Schematic of IFNγ ELISpot assay. For screening of CD8+ T cell and CD4+ T cell responses, CD8+ or CD4+ T cells were isolated from the spleens of tumor bearing mice and co-cultured with total splenoyctes from a healthy tumor-naïve mouse. **(D)** Representative IFNγ ELISpot images of assay peformed with CD8+ T cells from the spleen of CIT6-bearing mice cultured with 160ug/ml of Brinp3 peptide CPAFLPCTV (top row) and negative control (no peptide, bottom row). **(E)** Quantification of average IFNγ ELISpot spot counts for CD8+ T cells from the spleen of CIT6-bearing mice cultured with different concentrations of Brinp3 peptide CPAFLPCTV. Each colored line represents data points from a different mouse (n=4 mice). **(F)** Representative IFNγ ELISpot images of assay peformed with CD8+ T cells from the spleen of CIT6-bearing mice cultured with 160ug/ml of Ttll4 peptide VPPSSLLPL (top row) and negative control (no peptide, bottom row). **(G)** Quantification of average IFNγ ELISpot spot counts for CD8+ T cells from the spleen of CIT6-bearing mice cultured with different concentrations of Ttll4 peptide VPPSSLLPL. Each colored line represents data points from a different mouse (n=4 mice). **(H)** Schematic of immune checkpoint inhibitor (ICI) treatment regimen and ELISpot analysis. **(I)** Average spot counts of ICI-treated splenocytes cultured with 80ug/ml of Ttll4 peptide vs negative control. Each line represents data points from a different mouse (n=6 mice). Statistical analysis in panels E, G, and I was done by paired t-test. * *p* < 0.05, ** *p* < 0.01. Error bars in panels E and G represent mean and standard deviation.

Candidate neoantigens identified through this pipeline were subsequently screened by *in vitro* IFNγ ELISPOT assay^22,23^. Whole splenocytes, CD8+ T cells, or CD4+ T cells from the spleen of CIT6 tumor-bearing mice were incubated with synthetic peptides corresponding to each candidate epitope sequence (**Fig. 4C**). We identified two peptides, CPAFLPCTV (resulting from a Brinp3 T436P mutation) and VPPSSLLPL (resulting from a Ttll4 Q454L mutation), which consistently elicited IFNγ production in sorted CD8+ T cells from CIT6-bearing mice in a peptide concentration-dependent manner (**Fig. 4D-G**). These peptides elicited no response from sorted CD4+ T cells (**Supp. Fig. 3A**). When the peptide assay was carried out with whole splenocytes (no prior sorting of CD8+ or CD4+ T cells), the dose-dependent response to VPPSSLLPL (Ttll4 Q454L mutation) was still observed, but the response to CPAFLPCTV (Brinp3 T436P mutation) was not detectable above background (**Supp. Fig. 3B**).

We next interrogated the effect of ICI therapy on the T cell response to VPPSSLLPL, focusing on VPPSSLLPL because its response was more clearly detected in whole splenocytes **(Supp. Fig. 3B)**. CIT6 tumors were treated with three doses of anti-PD-1 and anti-CTLA-4 combination therapy (**Fig. 4H**), and an IFNγ ELISPOT assay was carried out on whole splenocytes harvested 3 days after the final dose of ICI. Splenocytes from ICI-treated mice produced a robust IFNγ response when incubated with VPPSSLLPL compared to no peptide **(Fig. 4I)**, indicating that response to the neoantigen VPPSSLLPL is prominent during immunotherapy response.

## Discussion

We have established the CIT lines as a novel series of *Hras*-driven cSCC tumor lines, and here present comprehensive histologic, genomic, and immune characterization of these 11 cell lines. The CIT lines carry genomic drivers relevant to human cSCC, as well as a moderate-to-high mutation burden^14,15^. We show that each CIT line exhibits a characteristic immune infiltrate, with individual cell lines ranging from immune hot to immune cold. Immune hot CIT lines CIT6 and CIT18 exhibit partial responses to an immunotherapy regimen of anti-CTLA-4 and anti-PD-1. We have also identified two neoantigens that elicit CD8+ T cell responses in the CIT6 cell line. Of note, while numerous neoantigens have been identified in tumor models syngeneic to C57BL/6^22–26^, there has been a notable dearth of neoantigen identification in other mouse strains including FVB, as well as a lack of neoantigens identified in cSCC models. The CIT lines, with the identification of two MHCI-restricted neoantigens in CIT6, thus provide a valuable new series of models for immunotherapy research in cSCC.

As noted above, the CIT cell lines are syngeneic to FVB mice. This stands in contrast to the majority of mouse models currently used for immunotherapy studies—across all cancer types—which are syngeneic to C57BL/6 mice. C57BL/6 mice are known to have a Th1-biased immune response^27^. While this is likely beneficial for tumor control—and C57BL/6 mice are widely known for their resistance to tumorigenesis^28,29^—it is not necessarily representative of the diversity of human immune responses. FVB mice, in contrast to C57BL/6, are known to exhibit Th2-biased immune responses^27,30^ and to be readily susceptible to tumorigenesis. While no single mouse model or mouse strain can represent the full spectrum of human immunity, increasing the diversity of mouse strains used to model anti-tumor immune responses is essential to capture human diversity and understand the full spectrum of barriers to immunotherapy success that are found in patients.

Finally, an additional feature of the CIT lines, having been derived from *K5CreER^T2^- Confetti* mice, is that cells can be labeled with fluorescent tags following exposure to Cre recombinase^31,32^. We have successfully labeled CIT lines *in vitro* using adenoviral Cre recombinase. While the *K5CreER^T2^* allele will also allow for labeling of K5+ (Keratin 5+) cells following tamoxifen treatment, we have found low expression of K5 in these poorly differentiated cell lines—as well as in DMBA/TPA tumors following progression to the carcinoma stage^31^—making this approach less effective. Introducing Cre and subsequently sorting cells enables the establishment of isogenic cell lines with each GFP, YFP, RFP or CFP^32^; while introducing Cre to activate labeling without sorting could allow for multiclonal lineage tracing.

Improving immunotherapy outcomes in advanced cSCC remains an unmet clinical need. The CIT panel of cutaneous SCC tumor lines provides an important new set of tools to better understand the mechanisms behind immunotherapy resistance in cSCC, and to preclinically model novel approaches to overcoming ICI resistance and improving cSCC patient outcomes.

## Materials and methods

### Animals

All animal experiments were approved by the by the University of Utah Office of Comparative Medicine (IACUC #22-02003) or by the University of California San Francisco Laboratory Animal Resource Center (IACUC #AN187679). This work complies with all the relevant ethical regulations regarding animal research. *K5CreER-Confetti FVB/N* mice were maintained in the Reeves Lab colony at the University of California San Francisco. Eight-week-old wild-type FVB/N mice were purchased from Jackson Laboratories. All experiments were performed on 8-16- week old mice. Any diversity in mouse age or sex was addressed by distributing age and sex cohorts evenly between different experimental arms.

### Carcinogenesis and cell line generation

To induce tumors, *K5CreER-Confetti FVB/N* mice were treated with 25 µg DMBA followed by TPA (200 µl of a 10^−4^ M solution in acetone) two times a week for 20 weeks. To generate cell lines, carcinomas were resected at a size of >1 cm in longest diameter, finely chopped, digested in DMEM containing Collagenase I and IV and DNase I for 45 minutes at 37 °C, washed with PBS, plated in DMEM plus 10% heat-inactivated FBS and L-glutamine, with penicillin, streptomycin, and amphotericin B, and passaged until cells stably grew in culture.

### In vivo experiments and immune checkpoint inhibitor treatment

A total of 1.25×10^5^ cells were injected subcutaneously into the dorsal flank of 8 to 16 week old male and female FVB/N mice. For baseline flow cytometric immune profiling and histological analysis (Fig. 1), tumors were harvested approximately 21 days post-engraftment, when tumors measured ∼1cm in diameter. For immune checkpoint inhibitor (ICI) experiments (Fig. 3), mice were enrolled in treatment 14 days post-tumor engraftement, when tumors grew to 25-100 mm^3^ in volume. Any mice with tumors <25 mm^3^ or >100 mm^3^ on the day(s) of enrollment were excluded from the experiment. Mice were treated with 200 µg anti-PD-1 (BioXCell Catalog# BE0146) and 200 µg anti-CTLA-4 (BioXCell Catalog# BE0131) or 200 µg isotype control (BioXCell Catalog# BE0089). Mice received two doses 3 days apart. Mice were harvested when tumors were ∼10mm in length, typically 17-21 days post-ICI enrollment and 31-35 days post-tumor engraftment, or when an ethical endpoint was reached.

### Flow cytometric immune profiling

Harvested tumors were finely chopped, digested in DMEM containing Collagenase IV (Sigma-Aldrich Catalog# 9001-12-1) and DNase I (Sigma-Aldrich Catalog# D5025-15KU) for 45 minutes at 37 °C, and filtered through 70um filters. Cells were counted using a Countess III, and 5 million cells were plated and stained with antibodies and Live/Dead NIR stain. Antibodies used for flow cytometric analyses are listed in Supp. Table 3. Flow cytometry analysis was performed on a Cytek Aurora.

### Histology

Harvested tumors were fixed in formalin for 24 hours, placed in cassettes (VWR Catalog# 18000), submerged in 70% ethanol, and submitted to ARUP Laboratories Research Histology Core Laboratory, where they were embedded in paraffin, cut into 4µm-thick slices, stained with hematoxylin and eosin (H&E) and mounted on slides. The 4µm slides were then analyzed by a board-certified pathologist, A.G.

### Cell line sequencing and analysis

DNA was extracted from cultured cells from each cell line and from tails of the mice in which each tumor originated using the Qiagen DNeasy Blood & Tissue DNA purification kit (Qiagen). Whole exome sequencing was performed using the Agilent Sure Select Mouse Exon capture kit, and sequenced to an average depth of 153X on an Illumina HiSeq X by MedGenome, Inc. in Redwood City, CA. Reads were trimmed with cutadapt (v3.5)^33^ and aligned to the mm10 mouse genome using BWA (v0.7.17)^34^. Reads were deduplicated using MarkDuplicates and recalibrated using BaseRecalibrator from GATK (v4.1)^35^. Mutations were called using MuTect2 (GATK v4.1)^35,36^ and Strelka2 (v2.9.10)^37^, using tails from the mice in which each tumor developed as the matched normal. Mutation calls were filtered using the following criteria: minimum read depth of 10 at the mutation position in both tumor and normal samples; minimum of 4 reads supporting the mutation call; for Strelka only, minimum quality QSS score of 25. Mutations that passed all filters with both callers were kept and were additionally filtered to remove any germline SNP detected in the panel of normals. Final mutation calls were annotated with Annovar^38^. Copy number calling was performed with CNVkit (v0.9.10)^39^, using all tails as a pooled normal. *Cdkn2a* deletions were confirmed by samtools (v1.16)^40^ mpileups of all reads aligning to the *Cdkn2a* locus (chr4:89,274,541-89,294,653). Exome sequencing data from CIT lines is available in SRA (accession number PRJNA1129114).

### Neoantigen peptide prediction

Mutations identified by exome sequencing were used to construct wild-type and mutant amino acid sequences of all 7mer, 8mer, and 9mer peptides that included each identified single nucleotide variant. The NetH2pan algorithm was used to predict binding affinity and percentile rank for each wild-type and mutant peptide sequence. This data was filtered using the following criteria: mutant peptide with a percentile rank <0.5, log affinity ratio >0 (affinity ratio defined as ratio of predicted binding affinity of mutant sequence dividesd by wild-type sequence), and a gene expression >300 RPM based on RNAseq of whole CIT6 and CIT9 tumors.

### ELISpot assays

Whole spleens were harvested from FVB mice and processed by crushing through a 70µm mesh filter, pre-wet with 1X PBS, using a syringe plunger and washed. Red blood cells were lysed for 5 minutes using ACK lysis buffer and splenocytes counted. Spleens from CIT6-bearing FVB mice were harvested 2-4 weeks after tumor inoculation. CD8+ T cells were isolated from whole spleens using EasySep^TM^ Mouse CD8+ T Cell Isolation Kits (STEMCELL Technologies Catalog# 19853) and CD4+ T cells were isolated from whole spleens using EasySep^TM^ Mouse CD4+ T Cell Isolation (STEMCELL Technologies Catalog# 19852). Interferon-gamma (IFNγ) ELISpots were performed over a three-day assay.

Multiscreen®HTS 96-well filtration plate(s) were coated with IFNγ capture antibody (clone AN-18, Catalog# 517902), diluted 1:500 in 1X PBS. Plates were wrapped in aluminum foil to equally distribute temperature and incubated overnight, or at least 18 hours, at 4°C. The following day plates were washed twice with 200µl RPMI-2 (RPMI with 2% FBS, 1% L-glutamate, 1% streptococcus and penicillin antibiotic solution, and 0.09% 2-Mercaptoethanol (BME)) and blocked for two hours with 200µl of RPMI-20 (RPMI with 20% FBS, 1% L-glutamate, 1% streptococcus and penicillin antibiotic solution, and 0.09% 2-Mercaptoethanol (BME)) before splenocytes or T cells were added. For assays where total splenocytes were used, 500,000 splenocytes were plated per well. For assays using CD8+ and CD4+ T cells, 20,000 cells per well were co-cultured with 500,000 splenocytes from a non-tumor-bearing mouse to provide a source of antigen presenting cells. Stimulation medium was prepared by diluting synthetic peptide in RPMI-10 to desired peptide concentration(s). All synthetic peptides used for these assays were ordered through GenScript. Peptides were reconstituted in DMSO or MilliQ water according to solubility testing provided by the manufacturer. Reconstituted peptides were aliquoted and frozen at −80°C. The negative control consisted of MilliQ water and RPMI-10.

Positive control contained 0.04% Phorbol 12-myristate 13-acetate (PMA) and 0.09% Ionomycin in RPMI-10. Plates were wrapped in aluminum foil and incubated undisturbed at 37°C and 5% CO2 for 44-48 hours before developing using biotinylated anti-IFNγ detection antibody (clone R4-6A2, Catalog# 505704), streptavidin-HRP enzyme conjugation (Thermofisher, Catalog# 405210), and AEC (3-amino-9-ethyl-carbazole) substrate. AEC substrate was prepared by dissolving 20 mg of AEC in 2 ml of N, N Dimethylforamamide (Sigma-Aldrich, Catalog# 227056), diluting in 0.1 M Acetate Solution, adding hydrogen peroxide and filtering with a 0.45µM filter. Spots were Imaged using the Agilent BioTek Cytation7 and analyzed using the Gen5 software.

### Statistical analysis

To determine significant statistical differences, Welch’s t-test was used when comparing 2 groups, and one-way ANOVA with Tukey post-hoc test was uses when comparing 3 or more groups. Paired analyses were performed by paired t-test. Hypothesis testing was performed using R open-source software (v4.4.2) and Prism software (GraphPad v10.4.2). Significant *p* values are marked as * p < 0.05, ** p < 0.01, *** p < 0.001, **** p < 0.0001. Studies were not blinded. Power calculations to determine appropriate cohort size were performed using Sample-Size software available online.

## Data availability

Exome sequencing of CIT lines is available in SRA (accession number PRJNA1129114).

## Acknowledgements

This work was supported by the UCSF Program for Breakthrough Biomedical Research (PBBR) Sandler Fellowship, US National Cancer Institute (NCI) grant R21CA264599, the Huntsman Cancer Institute Cancer Center Support Grant P30CA040214, the American Cancer Society IRG-21-131-01, Five For The Fight, The Cancer League, and the V Foundation (V2024-002). G.S. is supported by the Institute of General Medical Sciences grant T32GM141848. The computational resources used for exome sequencing analysis were partially funded by the NIH Shared Instrumentation Grant 1S10OD021644-01A1. We thank the University of Utah Flow Cytometry Core; the Associated Regional and University Pathologists, Inc. (ARUP Laboratories) Research Histology Core; the University of Utah Center for High Performance Computing (CHPC); the Huntsman Cancer Institute Precision Cancer Models (PCM) Shared Resource (supported by P30CA042014), in particular D. Lum and W. Sun; the Immune Oncology Network (ION), in particular E. Cortes-Sanchez; and the UCSF Parnassus Flow CoLab. We also thank A. Lai for help establishing the ELISPOT methods. We thank and all members of the Reeves Lab for thoughtful input during the development of this story. Some figures were created with Biorender.com.

## Author contributions

M.T., R.L., L.L., A.B., and M.Q.R. designed the project and experiments. R.L., S.H., M.T., L.L. and M.Q.R. led experiment implementation and data analysis. A.B. designed and carried out the neoantigen identification experiments. S.H. and G.S. carried out the immune checkpoint inhibitor experiments, with support from R.L. Board-certified pathologist A.G. provided histology review. R.L., M.T., D.C.D., and M.Q.R. wrote the manuscript, with input from other co-authors. P.C., K.N., D.S., C.M.L. and M.G. assisted with experiments.

---

**Supp. Fig. 1.**
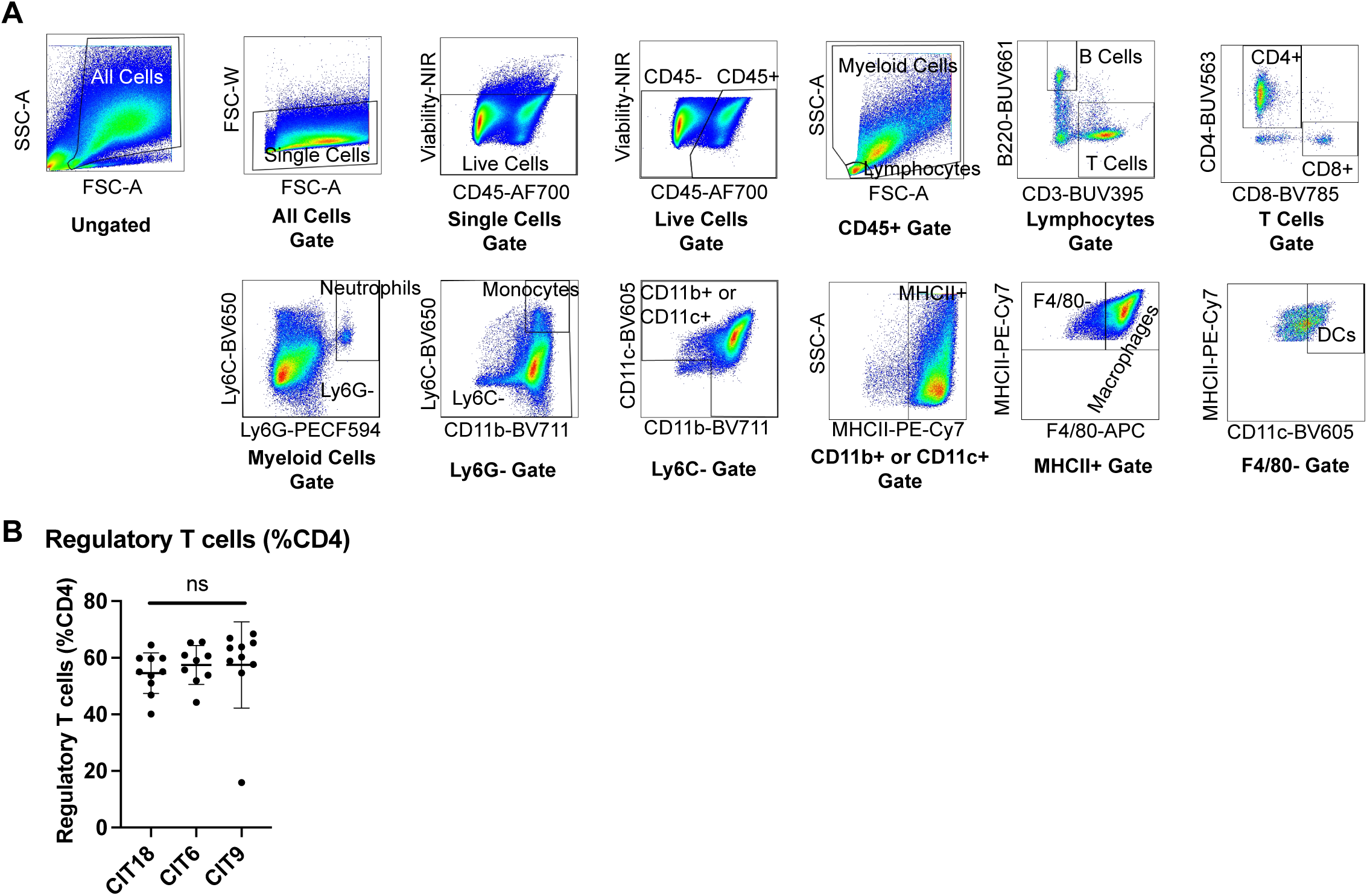
Flow cytometry gating strategy and regulatory T cell infiltration in CIT tumors. **(A**) Flow cytometric gating strategy of main immune cell types. Immune analysis of a CIT6 tumor is shown as representative data. **(B)** Abundance of Foxp3+CD25+ regulatory T cells (Tregs) as percentage of CD4+ T cells in CIT18, CIT6 and CIT9 tumors (n=9-10 mice per group). Statistical analysis by One-way ANOVA with Tukey post-hoc test.

**Supp. Fig. 2.**
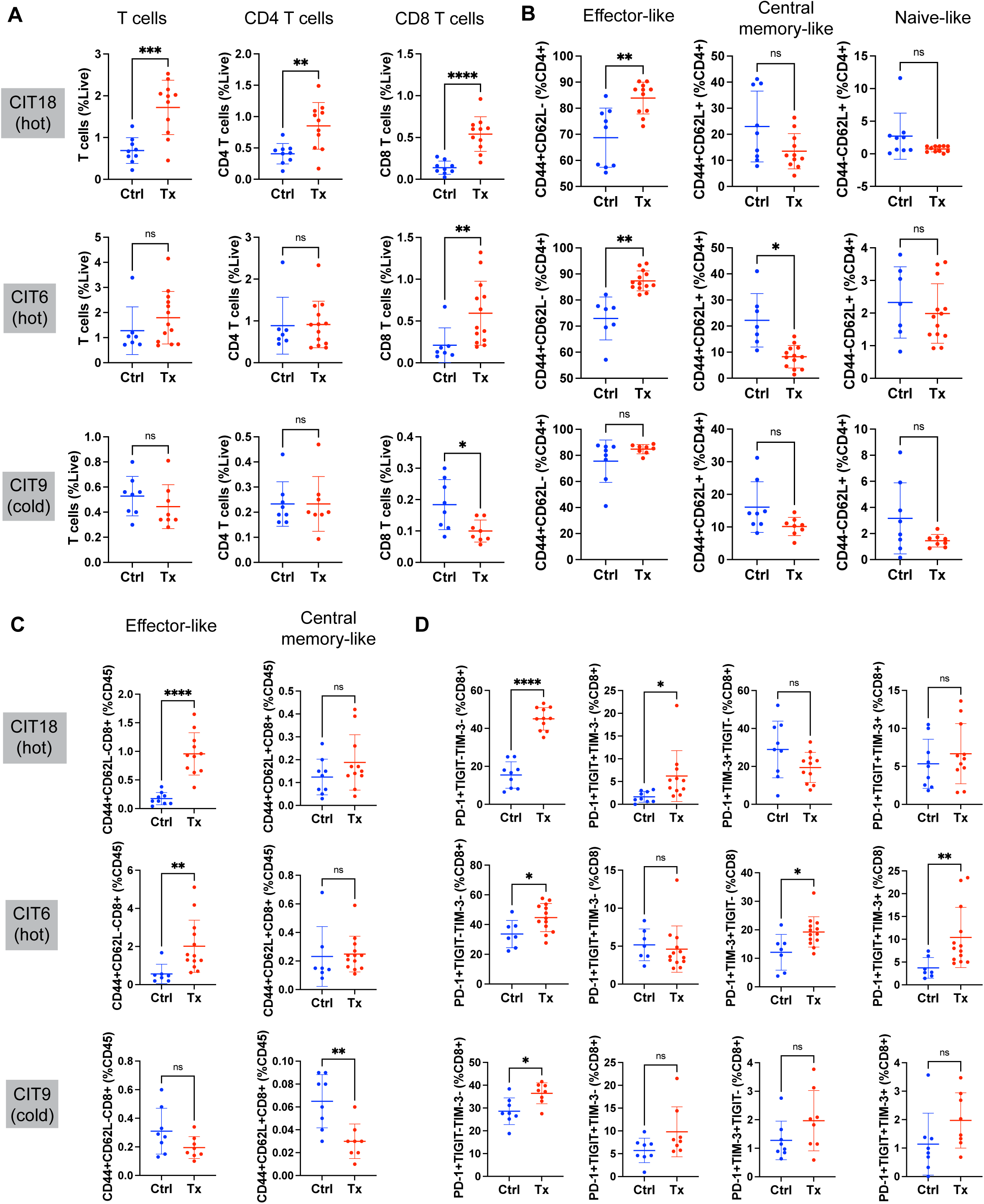
Tumor-infiltrating T cell effector activity after treatment with combined anti-PD-1 and anti-CTLA-4 immune checkpoint inhibitor (ICI) therapy. **(A)** Total T cell, CD4+ T cell, and CD8+ T cell infiltration of CIT18, CIT6 and CIT9 as percentage of all live cells, including tumor and stromal cells, for control-treated (“Ctrl”) or ICI-treated (“Tx”) tumors. **(B)** Abundance of effector subsets of CD4+ T cells defined by the combinatorial expressions of CD44 and CD62L in CIT18, CIT6 and CIT9. **(C)** Infiltration of CD44+CD62L- “effector-like” CD8+ T cells and CD44+CD62L+ “central memory-like” CD8+ T cells as percentage of all CD45+ immune cells. **(D)** Abundance of CD8+ PD-1+ T cells expressing specific combinations of TIGIT and Tim3 exhaustion markers in CIT18, CIT6 and CIT9 tumors. All statistical analyses performed with Welch’s t-test, n=7-13 mice per group. * *p* < 0.05, ** *p* < 0.01, *** *p* < 0.001, **** *p* < 0.0001.

**Supp. Fig. 3.**
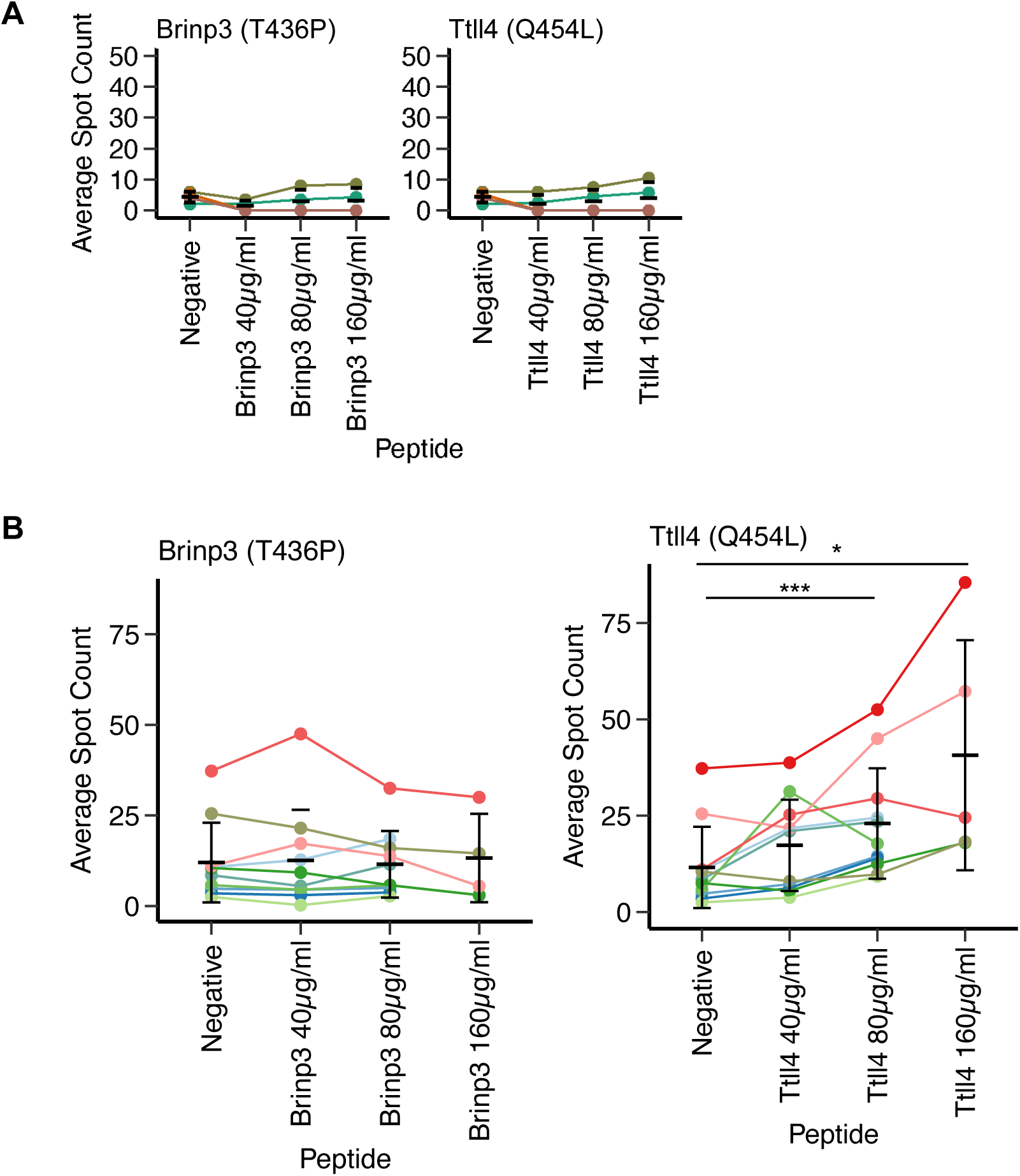
CIT6 ELISpot analysis of unfractionated splenocytes and CD4+ splenocytes. **(A)** Spot counts from ELISpot assays performed with CD4+ T cells isolated from the spleens of CIT6-bearing mice and cultured with splenocytes from tumor-naïve mice loaded with increasing concentrations of Brinp3 and Ttll4 peptides. Each colored line represents data points from a different mouse (n=4 mice). **(B)** Spot counts from ELISpot assays performed with total splenocytes from CIT6-bearing mice cultured with increasing concentrations of Brinp3 and Ttll4 peptides. Each colored line represents data points from a different mouse (n=10-11 mice). All statistical analyses performed with paired t-test. * *p* < 0.05, *** *p* < 0.001. Error bars represent mean and standard deviation.

**Supp. Table 1:** Number of total and non-synonymous mutations per CIT cell line. Number of total mutations, mutations per megabase, and non-synonymous mutations based on exome sequencing of CIT cell lines.

| **Cell Line** | **Total Mutations** | **Total Mutations per MB** | **Non-Synonymous Mutations** |
| --- | --- | --- | --- |
| CIT18 | 503 | 14.92 | 170 |
| CIT16 | 259 | 7.76 | 86 |
| CIT6 | 703 | 20.95 | 256 |
| CIT17 | 258 | 7.74 | 74 |
| CIT19 | 179 | 5.36 | 56 |
| CIT20 | 324 | 9.66 | 113 |
| CIT21 | 309 | 9.32 | 111 |
| CIT22 | 264 | 7.91 | 94 |
| CIT23 | 468 | 14.13 | 176 |
| CIT24 | 174 | 5.22 | 46 |
| CIT9 | 252 | 7.58 | 102 |

**Supp. Table 2:** List of filtered peptide candidates for CIT6 neoantigens. List of all neoantigen peptide candidates that met the criteria specified in Fig. 4A. Percentile ranked based on NetH2pan algorithm.

| Gene | Amino Acid Change | Peptide | Allele | Percentile Rank |
| --- | --- | --- | --- | --- |
| Slc39a10 | S286I | LPEHIGHEL | Dq | 0.0225 |
| Ttll4 | Q454L | VPPSSLLPL | Dq | 0.1367 |
| Brinp3 | T436P | CPAFLPCTV | Dq | 0.2119 |
| Cep76 | P158A | VPCACEADF | Dq | 0.4240 |
| Atp9a | I216F | EPNIDFHNF | Dq | 0.4662 |
| Fam83f | Q4L | VPALPTESL | Dq | 0.1392 |
| Emc1 | H307L | SHYALLHYL | Dq | 0.2106 |
| Cnot10 | A543P | VPLALGDNL | Dq | 0.0063 |
| Fam13b | D766H | QEKLARHL | Kq | 0.1266 |
| Setbp1 | N1160Y | HDYLSGLF | Kq | 0.2688 |
| Pcdh18 | M631I | NEENIFII | Kq | 0.1341 |
| Cpt1c | Q29H | HEIYLCAL | Kq | 0.3203 |
| Trip4 | Q423L | HQHQLRIL | Kq | 0.3234 |

**Supp. Table 3:** List of antibodies used for flow cytometry analysis. All antibodies, clone number, and conjugated fluorophores for antibodies used for flow cytometry.

| Target | Fluorophore | Clone | Dilution | Manufacturer | Catalog # |
| --- | --- | --- | --- | --- | --- |
| B220 | BUV661 | RA3-6B2 | 1:200 | BD Biosciences | 612972 |
| CD11b | BV711 | M1/70 | 1:100 | BioLegend | 101242 |
| CD11c | BV605 | N418 | 1:100 | BioLegend | 117334 |
| CD16/CD32<br>(FcR Block) | N/A | 2.4G2 | 1:167 | Tonbo | 70-0161-U500 |
| CD3e | BUV395 | 145-2C11 | 1:62.5 | BD Biosciences | 563565 |
| CD4 | BUV563 | GK1.5 | 1:200 | BD Biosciences | 612923 |
| CD44 | BV785 | 1M7 | 1:200 | BioLegend | 103059 |
| CD45 | AlexaFluor700 | 30-F11 | 1:400 | BioLegend | 103128 |
| CD62L | FITC | MEL-14 | 1:200 | Invitrogen | 11-0621-81 |
| CD8a | BV570 | 53-6.7 | 1:200 | BioLegend | 100739 |
| CD8a | BV785 | 53-6.7 | 1:200 | BioLegend | 100750 |
| F4/80 | APC | BM8 | 1:100 | BioLegend | 123116 |
| FoxP3 | PE-Cy7 | FJK-16s | 1:50 | eBioscience | 25-5773-82 |
| Live/Dead | NIR | N/A | 1:1000 | Invitrogen | L34976A |
| Ly6C | BV650 | HK1.4 | 1:200 | BioLegend | 128049 |
| Ly6G | PE-CF594 | 1A8 | 1:200 | BD Biosciences | 562700 |
| MHCII | PE-Cy7 | M5/114.15.2 | 1:400 | BioLegend | 107630 |
| PD-1 | BV605 | 29F.1A12 | 1:100 | BD Biosciences | 135220 |
| TIGIT | BV421 | 1G9 | 1:100 | BioLegend | 142111 |
| Tim3 | APC | RMT3-23 | 1:100 | BioLegend | 119705 |

